# Track Display Jockey (trackDJ): a user-friendly R package for visualization of epigenomic data

**DOI:** 10.64898/2026.04.15.718328

**Authors:** Neha V. Bokil, David C. Page

**Affiliations:** Whitehead Institute, Cambridge, MA 02142, USA; Department of Biology, Massachusetts Institute of Technology, Cambridge, MA 02139, USA; Howard Hughes Medical Institute, Whitehead Institute, Cambridge, MA 02142, USA

## Abstract

**Background:** Visualization of epigenomic data such as coverage tracks, peak calls, and chromatin interactions is a critical task in genomic data analysis. Although genome browsers such as the Integrative Genomics Viewer (IGV) and the UCSC Genome Browser permit user-friendly exploration of genomic tracks, they are not optimized for fully programmatic and reproducible generation of publication-quality figures. In contrast, existing programmatic tools lack a user-friendly interface and require extensive configuration.

**Results:** We present **trackDJ (Track Display Jockey)**, an R package for visualization of epigenomic data. *trackDJ* prioritizes usability by favoring convention over configuration; it provides high-level plotting functions with sensible defaults, aiming to allow users with minimal programming experience to generate clear, publication-quality figures with relatively little coding. Within a unified plotting framework, users can stack and align multiple data types, including coverage tracks, peak annotations, chromatin loops, and gene annotations. *trackDJ* allows users to select plotted genomic regions by coordinates or by gene name, enabling rapid visualization without knowledge of precise locus boundaries.

**Conclusions:** *trackDJ* complements existing epigenomics toolkits by providing a user-friendly, reproducible alternative to interactive genome browsers for epigenomic visualization. By enabling scripted generation of clean, customizable genomic illustrations, *trackDJ* readily integrates into R-based analysis workflows to streamline the creation of publication-quality figures.

## Background

High-throughput sequencing assays such as ChIP-seq [1,2], ATAC-seq [3], CUT&RUN [4], bisulfite sequencing [5], and Hi-C [6] are widely used to study histone marks, chromatin accessibility, transcription factor binding, DNA methylation, and genome organization.

Effective visualization of these data across genomic loci is essential for quality control, hypothesis generation, data interpretation, and communication of results. Typical visualizations integrate multiple data types, including signal coverage, chromatin interactions, and gene annotations.

Interactive genome browsers such as the Integrative Genomics Viewer (IGV) [7] and the UCSC Genome Browser [8] provide powerful exploratory visualization and do not require programming expertise. However, producing publication-ready figures using these tools often involves manual, labor-intensive steps to configure tracks, genomic regions, colors, and labels. This process is often difficult to integrate into analysis pipelines, and it is challenging and time-consuming to reproduce figures exactly.

Within the R ecosystem, extensive infrastructure exists for genomic data import and manipulation. However, while several R packages support genomic visualization (e.g., *Gviz* [9], *ggbio* [10], *ggcoverage* [11]), these tools have a steep learning curve, require substantial configuration, may not integrate seamlessly with *ggplot2* [12]-based workflows, and lack convenient support for common data types such as coverage tracks and chromatin loops.

Constructing clear, multi-track genomic figures with these tools often requires substantial plotting expertise and laborious adjustments in secondary design software such as Adobe Illustrator. This creates a barrier for users with limited programming backgrounds and can lead to inconsistent figure styling across analyses.

To address these challenges, we developed *trackDJ (Track Display Jockey)*, an R package designed to provide an intuitive, programmatic solution for generating high-quality genome browser-style visualizations. By leveraging the widely-used *ggplot2* framework and *rtracklayer* [13] for data import, *trackDJ* integrates seamlessly with existing Bioconductor workflows while minimizing the amount of code required to produce figures. *trackDJ* emphasizes ease of use, sensible defaults, and clean figure output without sacrificing customization, enabling researchers to easily and efficiently generate reproducible visualizations across multiple experimental conditions.

## Implementation

### Software architecture

*trackDJ* is implemented in R and builds upon established Bioconductor infrastructure. The package architecture consists of three core components: data import, plot generation, and track assembly (Figure 1).

**Figure 1:**
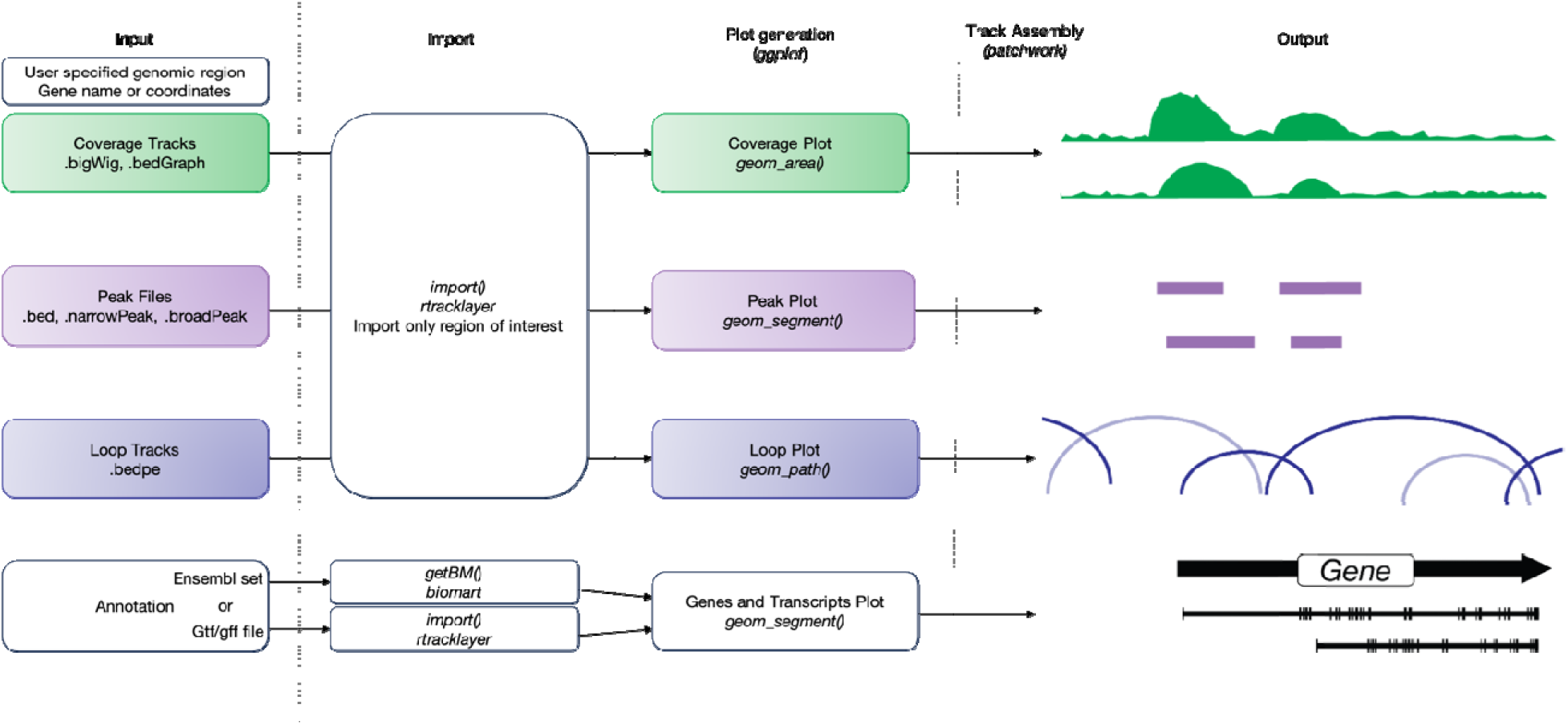
Package architecture of *trackDJ. trackDJ* imports annotations, coverage tracks, peak files, and loop tracks. Plots are generated using *ggplot2* and assembled with *patchwork* to output a stacked figure at a specified genomic locus.

Data import leverages *rtracklayer* [13] to read tracks from the standard genomic formats bigWig, bedGraph, bed, and bedpe. This approach ensures compatibility with output from popular analysis pipelines such as *deepTools* [14] and *BEDtools* [15].

For each track type, *trackDJ* generates a separate *ggplot* object. To ensure that tracks of the same type will have consistent sizing, *trackDJ* uses faceting while generating these plots. *trackDJ* then generates a patchwork [16] object that combines all tracks into a single figure. The output includes both the final figure and the underlying *ggplot* objects. If desired, *trackDJ* can use these *ggplot* objects to change the order in which tracks are displayed.

### Genomic annotations

*trackDJ* integrates with Ensembl through the *biomaRt* [17] package to retrieve gene and transcript annotations. Users can specify the genome assembly and plot transcripts if desired. Alternatively, to support genome builds not available in Ensembl, *trackDJ* allows users to provide genomic annotations as gtf/gff3 files or *GRanges* [18] objects.

To specify the region to be displayed, *trackDJ* supports both gene-centric and coordinate-based approaches. Users can query by gene symbol or provide explicit genomic coordinates. Setting `gene_symbol` to “ensembl_gene_id” allows users to input stable Ensembl gene IDs to select genes. When a gene identifier is provided, *trackDJ* automatically retrieves the gene coordinates using the selected annotation resource. Additionally, users can specify the number of base pairs upstream and/or downstream of a selected gene to plot, expanding the viewing window without explicit genomic coordinates.

### Emphasis on usability

*trackDJ* was designed with usability as a primary consideration. Common visualization tasks are executed through high-level functions that require minimal configuration. Sensible defaults are provided for track spacing, scaling, resolution, and aesthetics, allowing users to generate clear figures with minimal coding. Additionally, to minimize clutter, *trackDJ* defaults to plotting only protein-coding gene loci rather than rendering all transcript isoforms. *trackDJ* generates a main figure as a *patchwork* object as well as each of the underlying *ggplot* objects. The latter provides easy access to the data used for plotting and allows users to apply standard *ggplot2* operations if further customization is desired. Users can utilize *ggsave* to export *trackDJ* plots to publication-quality vector formats such as SVG and PDF, or raster formats such as PNG and TIFF.

### Track Customization

By default, *trackDJ* generates black-and-white figures and labels tracks according to their type and input order (e.g., coverage tracks: *Cov_1*, *Cov_2*; peak tracks: *Peaks_1*, *Peaks_2*; loop tracks: *Loops_1*, *Loops_2*). Users can customize each track in a *trackDJ* plot through a consistent set of parameters. All types of tracks accept specifications for color and labels, enabling clear identification of different experimental conditions. For coverage tracks, users can specify maximum and minimum values for the y-axis, and there is an option to select a logarithmic scale instead of the default linear scale. Peak track customizations include the ability to label specific peaks, to color specific peaks differently from others, and indicate strand, if applicable. For loop tracks, users can specify line thickness and loop orientation. Additionally, users can provide a score threshold such that loops with a lower score will be shaded more lightly.

Under default parameters, *trackDJ* plots only gene models of protein-coding loci, but users have the option to include non-coding genes by setting `includeNPC` to `TRUE`. Users can plot transcripts by setting `includeTranscripts` to `TRUE`. When plotting transcript models, users can elect to visualize all transcript isoforms in an annotation or filter them to include only a subset. Filtering options include plotting only the canonical transcript, specifying a list of transcripts to plot, or removing low-confidence annotations based on the `transcript_support_level` attribute (in Ensembl) and/or APPRIS classification [19].

Users can also specify other attributes they wish to use for filtering; in this case, *trackDJ* will only plot transcripts with a non-empty field for the given attribute.

## Results

### Programmatic creation of browser-style views

We demonstrate *trackDJ* functionality through representative use cases spanning common epigenomic applications. All data used in these demonstrations are available for download through the ENCODE database [20] (Supplementary Table 1).

All types of tracks can be plotted simultaneously with the `plot_genomic_tracks` function. Minimally, users need only supply a genomic locus and the filepaths to each of their tracks (Figure 2A):

**Figure 2:**
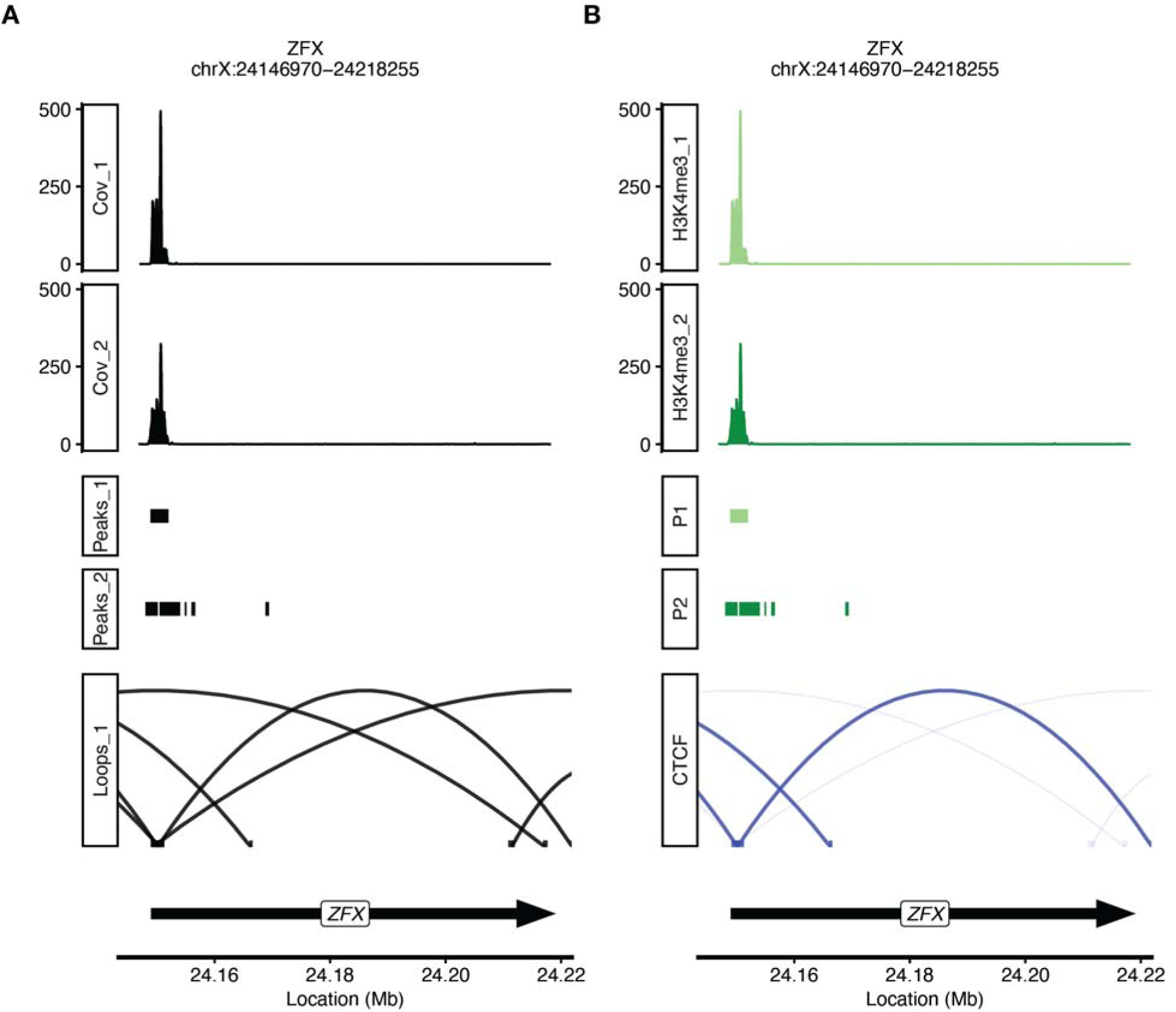
H3K4me3 ChIP-seq data coverage tracks and peaks followed by CTCF ChIA-PET data across a gene of interest (*ZFX*). A: *trackDJ* figure with default settings. B: *trackDJ* figure after specifying labels and colors of each track and shading for loops.

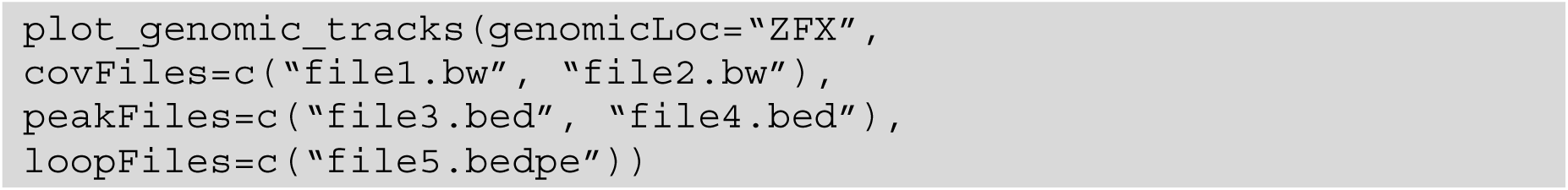

Users can easily customize track labels and colors to provide clarity (Figure 2B):

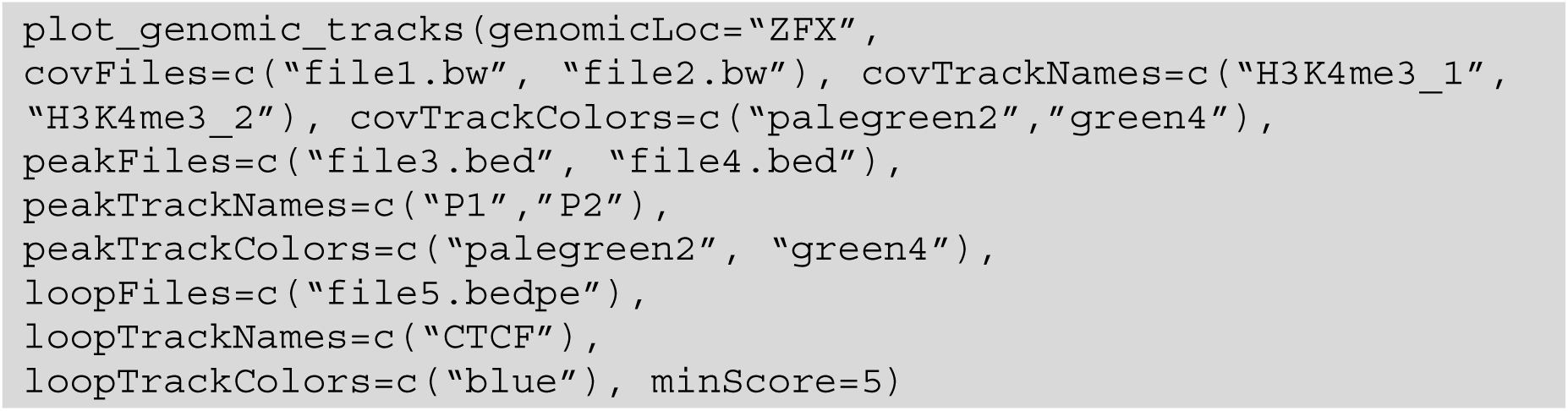

Instead of a gene name, users can provide a set of genomic coordinates (Figure 3).

**Figure 3:**
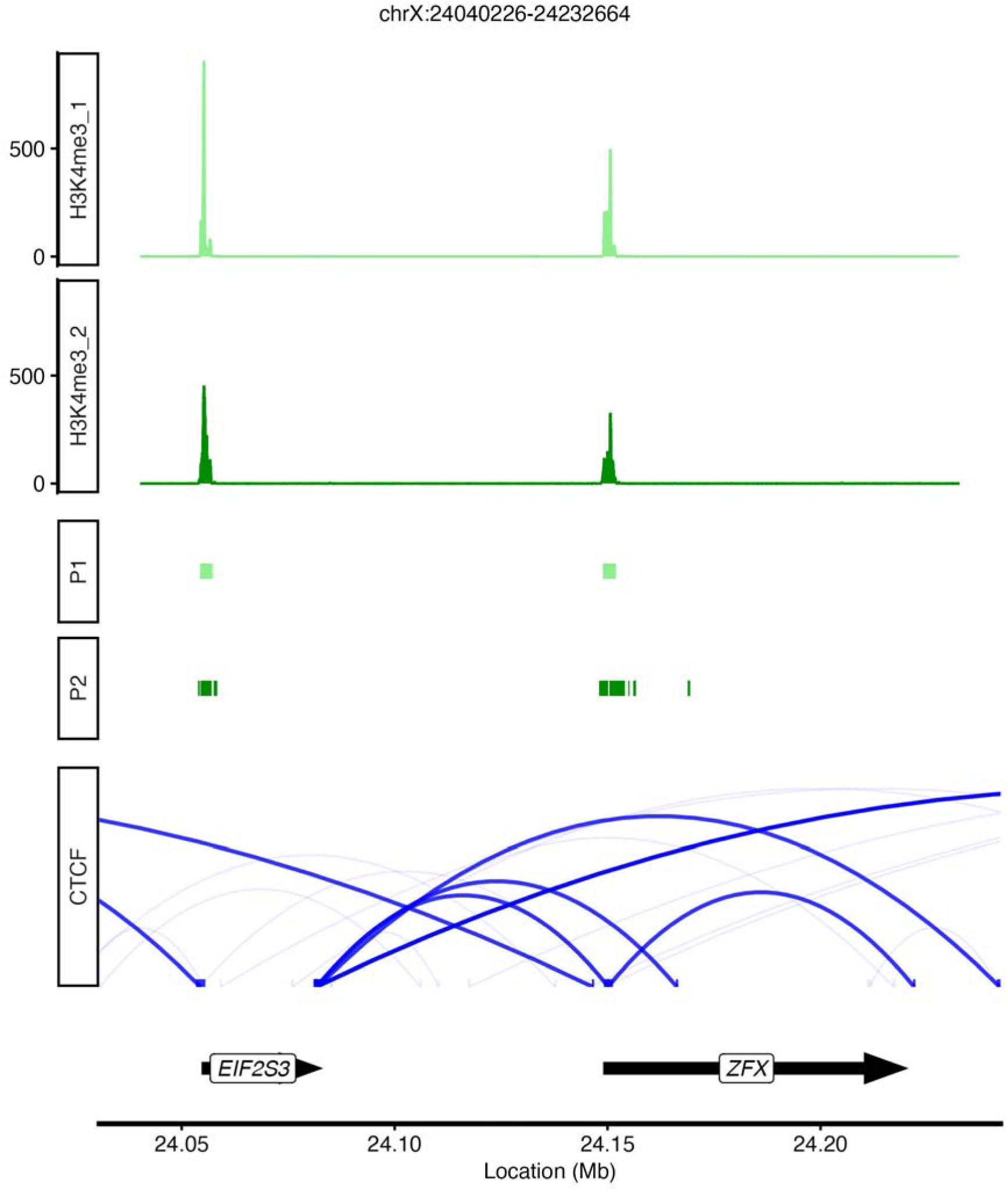
H3K4me3 ChIP-seq data (green) and CTCF ChIA-PET data (blue) across a genomic region of interest specified by coordinates.

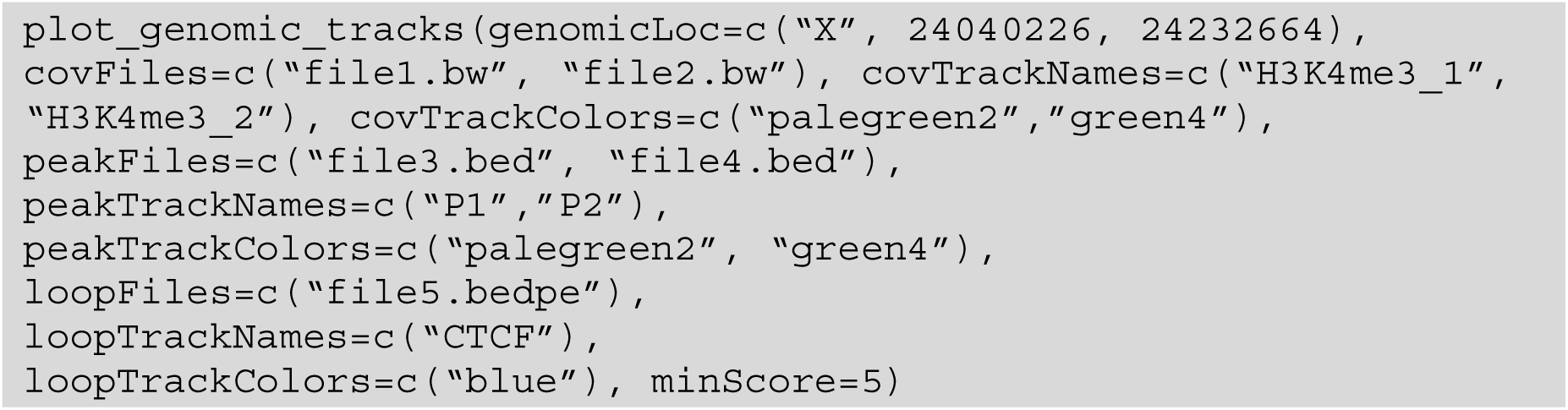

The `plot_genomic_tracks` function features a number of options to further customize images. Users can specify their desired order of track types. Depending on the application, users may want to show transcript isoforms alongside gene models, especially when epigenomic marks differ between isoforms of the same gene. Putting coverage tracks on a log scale can make subtle but biologically meaningful differences visible, particularly when signal ranges span several orders of magnitude. Highlighting specific peaks in a different color allows users to quickly distinguish regions of interest. If gene models are at the top of the plot rather than the bottom, flipping the orientation of loop tracks can make spatial relationships clearer. Users can make each of these adjustments simultaneously (Figure 4).

**Figure 4:**
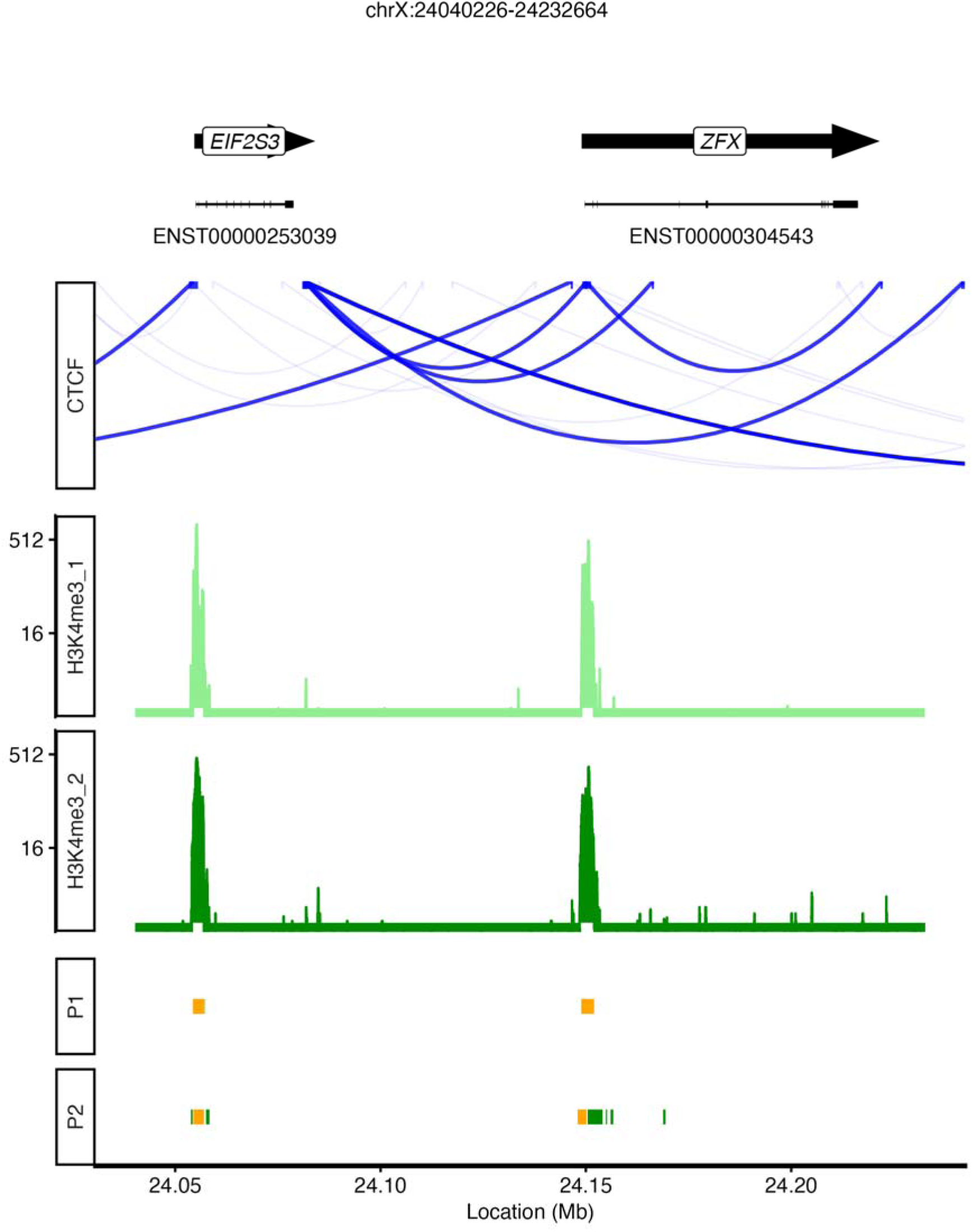
H3K4me3 ChIP-seq data (green) and CTCF ChIA-PET (blue) across the same region of interest shown in Figure 3, with modifications. This includes: plotting canonical transcripts, flipping loop orientation, putting coverage tracks on a logarithmic scale, coloring peaks of interest differently (orange), and changing the order of tracks.

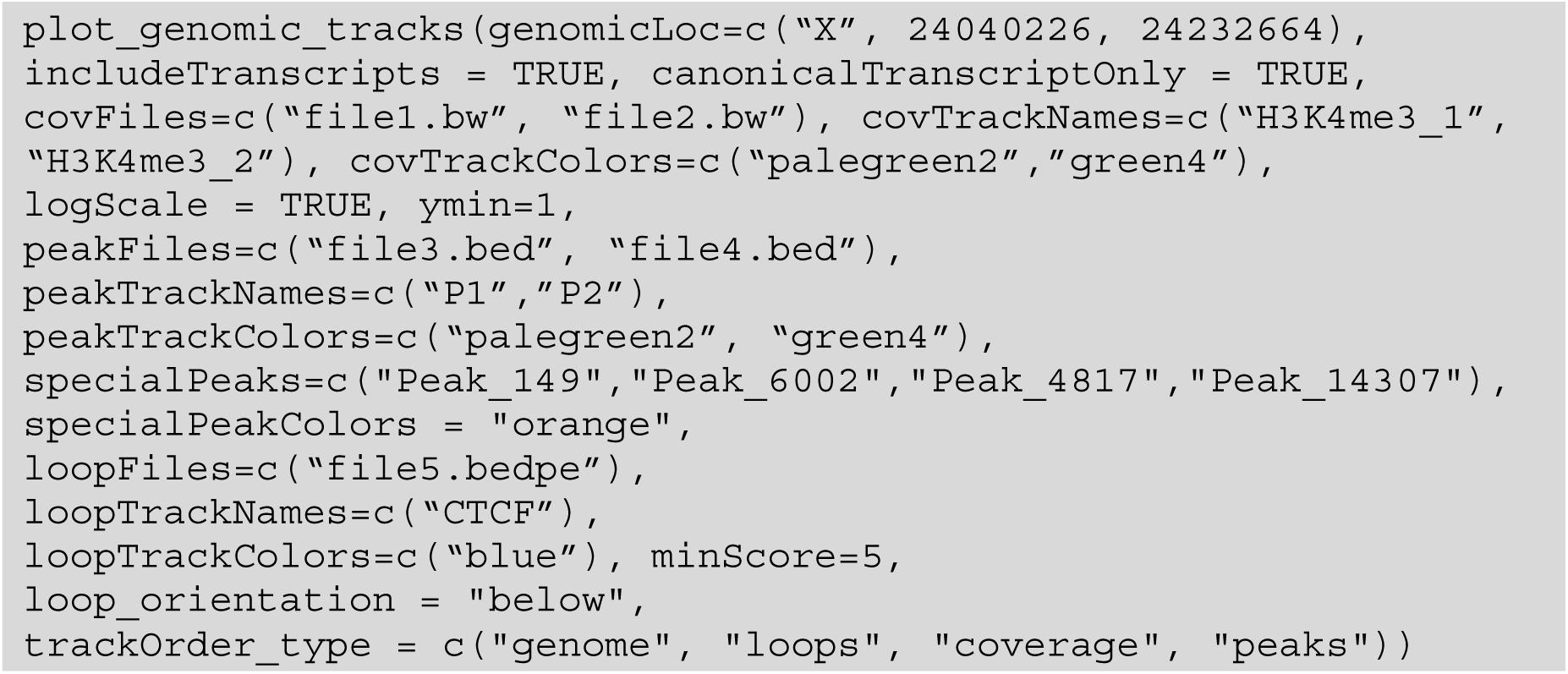

The default annotation in *trackDJ* is the Ensembl set hsapiens_gene_ensembl, but users can specify other annotations instead. For example, users can provide another Ensembl set such as the mouse annotation mmusculus_gene_ensembl (Supplementary Figure S1). *trackDJ* can also handle custom annotation files in gtf and gff3 format, allowing users to plot data from infrequently studied organisms such as the vicuña (*Lama vicugna*) [21] (Supplementary Figure S2).

Because `plot_genomic_tracks` plots all coverage tracks on the same scale, users must generate a different plot for each scale being used. To combine these plots into one, users can pass all `plot_genomic_tracks` outputs into the `trackDJ` function together with the desired track order. Unlike `plot_genomic_tracks`, which groups tracks of the same type together, `trackDJ` “mixes” the tracks so that they can all be displayed in the specified order in one figure (Figure 5).

**Figure 5:**
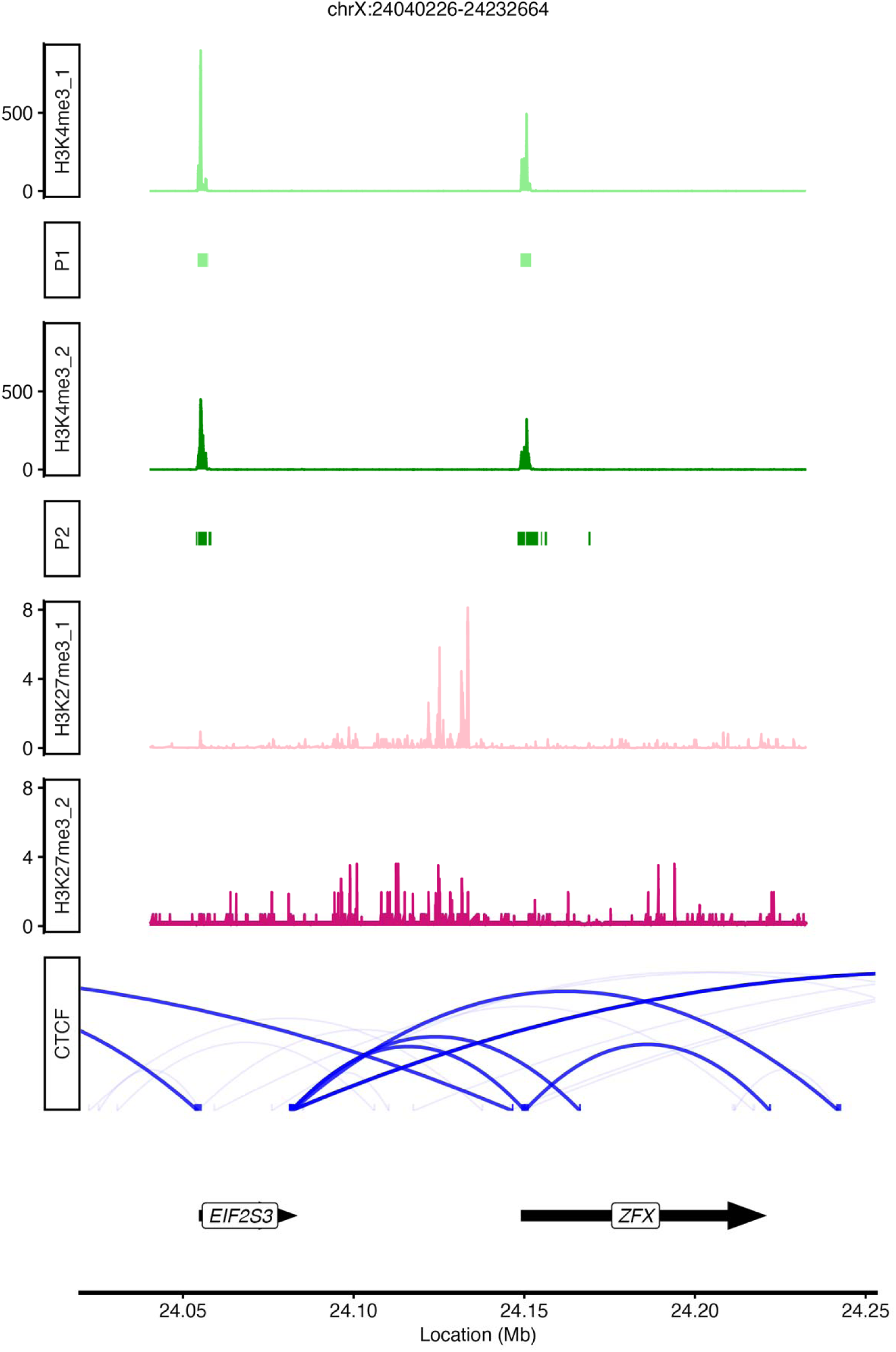
H3K4me3 ChIP-seq data (green), H3K27me3 ChIP-seq data (pink), and CTCF ChIA-PET (blue) across the same region of interest shown in Figures 3 and 4. Coverage tracks have different scales and tracks have been reordered such that H3K4me3 peaks are placed under their respective coverage tracks.

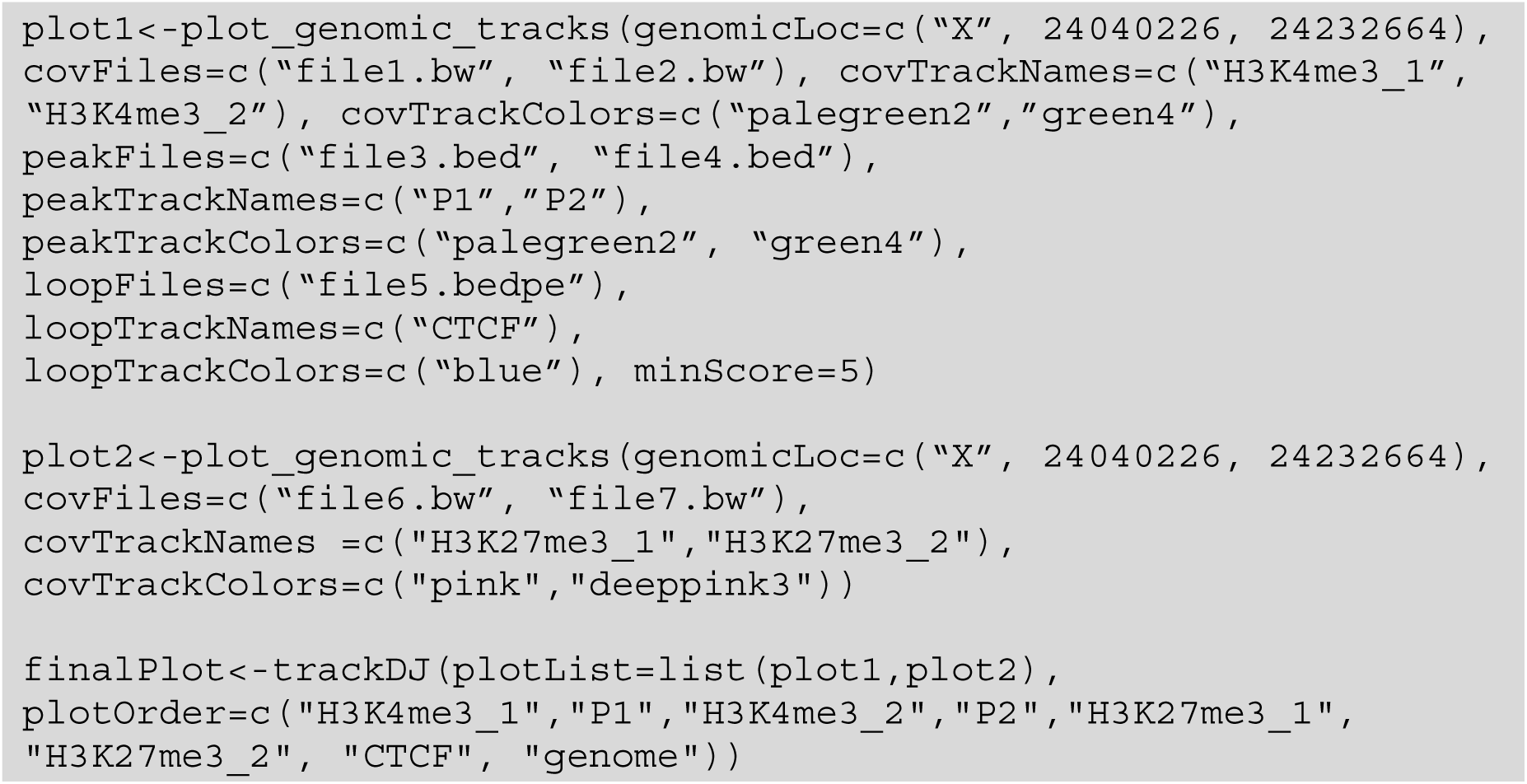

### Performance characteristics

Benchmarks were run using R version 4.5.3 on a 64-bit macOS system (macOS 15.6.1, ARM64/Apple Silicon). All benchmarks were run in a single-threaded configuration, with OpenMP threads restricted via OMP_NUM_THREADS and BLAS threads restricted via the Accelerate framework’s VECLIB_MAXIMUM_THREADS environment variable. Time was measured using the `system.time` function in R, and memory usage was monitored using the *peakRAM* R package [22] version 1.0.2. We ran `plot_genomic_tracks` 100 times, each time randomly selecting one out of four possible regions of different lengths: a 2.5 kb region specified by coordinates (chrX: 24148500-24151000), a 24.4kb gene (*LTBP3*), a 71.3kb gene (*ZFX*), or a 244kb gene (*KDM6A*). Each plot generated consisted of four coverage tracks (file1.bw, file2.bw, file6.bw, file7.bw), two peak tracks (file3.bed, file4.bed), and one loop track (file5.bedpe), for a total of seven input files. For the annotation, we generated an Ensembl mart object once prior to timing and supplied it to all 100 replicates, avoiding the overhead of repeatedly re-establishing a connection to Ensembl. However, gene coordinate information was not cached across replicates, so each of the 100 runs still issued an annotation query against this mart. Reported runtimes therefore reflect the per-call cost of annotation retrieval but exclude the one-time cost of mart initialization.

In our benchmark, *trackDJ* generated each figure in a median time of 1.77 seconds (Supplementary Figure S3A) on the hardware described above. Peak memory usage did not exceed 0.02 GB in any run (Supplementary Figure S3B). Because *rtracklayer* imports only the requested genomic intervals rather than whole files, memory use is expected to remain low for region-level plots.

### Comparison with existing genomic visualization tools

Several R packages provide functionality for visualizing genomic data, including *Gviz* [9], *ggbio*[10], and *ggcoverage* [11]. *Gviz* and *ggbio* are designed as general-purpose frameworks that offer extensive flexibility across a wide range of genomic data types, whereas *ggcoverage* is focused on visualizing and annotating sequencing coverage. However, these tools are not optimized specifically for producing publication-style epigenomic track figures with minimal configuration.

*Gviz* implements a comprehensive system of genomic track objects based on grid graphics, enabling detailed customization of genomic visualizations. While this approach is powerful, it typically requires explicit construction and coordination of multiple track objects and familiarity with a complex S4 class hierarchy. For routine epigenomic visualization tasks, such as stacking signal coverage tracks with peak annotations and gene models, this can result in verbose code and substantial configuration overhead. *trackDJ* adopts a more opinionated design, providing streamlined functions tailored to commonly used epigenomic data. Unlike *Gviz*, *trackDJ* produces *ggplot2* objects by default and is designed to integrate seamlessly into modern tidyverse-based workflows.

*ggbio* extends the grammar of graphics to genomic data and offers powerful abstractions for coordinate transformations and feature visualization. However, this abstraction can introduce a steep learning curve for users seeking to reproduce genome browser–style figures. Effective use of *ggbio* often requires users to directly manage genomic coordinate transformations and plot structure. *trackDJ* instead emphasizes a higher-level interface that maps directly to epigenomic visualization concepts, such as coverage tracks, peak tracks, chromatin loops, and gene annotations.

*ggcoverage* is a *ggplot2*-based package for visualizing and annotating sequencing coverage across a range of NGS data types. It provides a set of dedicated layer functions for common genomic features, including `geom_gene()` and `geom_transcript()` for gene models; `geom_peak()` for peak annotation; `geom_link()` and `geom_tad()` for chromatin interaction data; and `geom_ideogram()` for chromosomal context. Once a base coverage plot is constructed from user-loaded track data, *ggcoverage* manages axis scaling and coordinate alignment across these layers internally. However, building a complete multi-track figure requires the user to have detailed knowledge of each layer function. Users must load track data and then additively compose multiple purpose-built `geom` layers using *ggplot2*’s ‘+’ operator. *trackDJ* instead wraps this composition into single high-level functions per figure type, reducing the number of steps required to go from raw data to a final figure. We also note a difference in the visual consistency of default output: representative multi-panel figures in the *ggcoverage* documentation and original publication [11] combine track types whose fonts, tick-label formatting, and color assignment vary from panel to panel, suggesting that a uniform appearance across an entire multi-track figure requires additional manual `theme()` and color-palette adjustments by the user. *trackDJ* instead applies a single, consistent default theme across all track types within a figure, so that multi-track outputs are visually uniform without further post-processing.

In contrast to each of these tools, *trackDJ* supports gene-name-based region specification, facilitating intuitive, gene-centric workflows. This feature is particularly useful for exploratory analyses and figure generation at specific genes of interest, with no prior knowledge of genomic coordinates necessary. These tools also differ from *trackDJ* in how track-level formatting is specified. In *Gviz*, color, fill, and label properties are set through each track class’s own display-parameter system, so achieving a particular look across different track types requires learning and coordinating separate sets of graphical parameters for each class. In *ggbio* and *ggcoverage*, formatting instead follows standard *ggplot2* conventions, meaning consistent colors and labels across tracks must be built and aligned as separate mapping calls or arguments applied to each layer individually. *trackDJ*, by contrast, utilizes simple, consistently named arguments for every track type—for example, `covTrackColors`, `peakTrackColors`, and `loopTrackColors` for setting colors, and their corresponding `covTrackNames`, `peakTrackNames`, and `loopTrackNames` for labels—so that a user can set the appearance of all types of tracks the same way without learning a different system for each data type.

Table 1 summarizes these differences, including a qualitative rating of each package’s configuration complexity, which we defined using three criteria: (i) the number of explicit function- or object-construction steps required to produce a standard multi-track epigenomic figure — a single high-level function call per track (low), explicit composition of generic grammar-of-graphics layers (moderate-high), or explicit instantiation and coordination of multiple S4 track-object classes (high); (ii) whether genomic coordinate transformations across tracks must be managed manually by the user rather than handled internally by the package (low if internal, moderate-high/high if manual); and (iii) whether dedicated epigenomic track types — coverage, peaks, chromatin loops, and gene models — are available as built-in defaults or must instead be assembled from general-purpose graphical primitives (low if built-in, high if not). Applying these criteria, *Gviz*’s requirement to explicitly construct and coordinate multiple S4 track objects places it in the high category; *ggbio*’s grammar-of-graphics abstraction, combined with its need for manual coordinate management, places it in the moderate-high category; *ggcoverage*, while ggplot2-native and providing dedicated layer functions for common epigenomic features with internally managed coordinate alignment, still requires users to additively assemble multiple layers per figure and to supply explicit chromosomal coordinates rather than gene names, placing it in the moderate category; and *trackDJ*’s single-call, epigenomics-specific interface places it in the low category. By prioritizing usability and minimal configuration for common epigenomic visualization tasks, *trackDJ* complements existing visualization frameworks while addressing a distinct and practical need for biologists hoping to interpret and share their findings (Table 1).

**Table 1:**
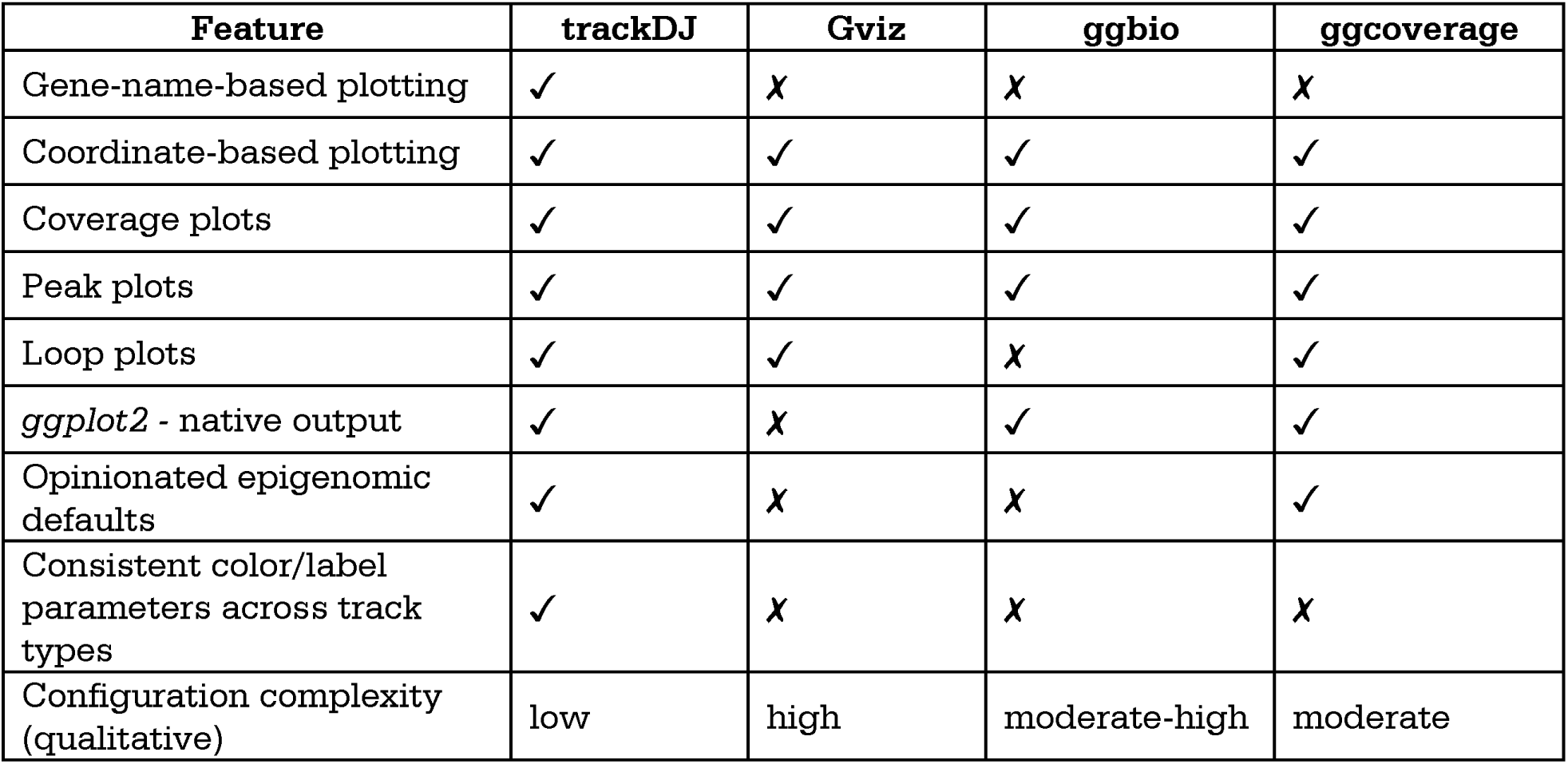
Features of *trackDJ* and other programmatic tools for plotting epigenomic data.

| **Feature** | **trackDJ** | **Gviz** | **ggbio** | **ggcoverage** |
|---|---|---|---|---|
| Gene-name-based plotting | ✓ | ✗ | ✗ | ✗ |
| Coordinate-based plotting | ✓ | ✓ | ✓ | ✓ |
| Coverage plots | ✓ | ✓ | ✓ | ✓ |
| Peak plots | ✓ | ✓ | ✓ | ✓ |
| Loop plots | ✓ | ✓ | ✗ | ✓ |
| *ggplot2* - native output | ✓ | ✗ | ✓ | ✓ |
| Opinionated epigenomic defaults | ✓ | ✗ | ✗ | ✓ |
| Consistent color/label parameters across track types | ✓ | ✗ | ✗ | ✗ |
| Configuration complexity (qualitative) | low | high | moderate-high | moderate |

## Discussion

*trackDJ* aims to enhance the epigenomics software ecosystem by providing a programmatic alternative to manual figure generation from genome browsers. While interactive browsers remain invaluable for exploratory analysis, the systematic generation of publication-quality figures requires automation and reproducibility that manual workflows cannot provide.

The design philosophy of *trackDJ* emphasizes usability without sacrificing functionality. In our benchmark, figure generation had a median runtime of 1.77 seconds and peak memory use did not exceed 0.02 GB for the tested configurations. *trackDJ* also enforces consistent alignment, spacing, themes, and annotation handling to simplify the creation of publication-ready figures. Transcript filtering options reduce visual clutter. Figures generated by *trackDJ* can be directly exported for use in manuscripts and reports.

A current limitation of *trackDJ* is the lack of direct remote (URL-based) file access. Because *rtracklayer* already supports importing remote genomic data [13], we opted to build on existing Bioconductor infrastructure rather than reimplement this functionality. Native remote file support could be added in future releases of *trackDJ* if it proves to be a common need.

By prioritizing ease of use and sensible defaults, *trackDJ* is designed to lower the barrier to epigenomic visualization for users with limited programming experience while still offering flexibility through its *ggplot*-based design. This combination aims to support reproducible research workflows and automated figure generation. The package’s modular architecture allows users to start with simple visualizations and progressively add complexity as needed.

## Conclusions

*trackDJ* aims to provide an accessible, programmatic solution for generating publication-quality visualizations of epigenomic data. By integrating with established R/Bioconductor tools and the *ggplot2* framework, *trackDJ* enables researchers to efficiently create figures without additional manual adjustment in genome browsers or graphics software. *trackDJ*’s high-level plotting functions allow figures to be created with minimal code. The package streamlines figure generation for manuscripts and presentations while maintaining the flexibility required for diverse experimental designs and visualization needs, facilitating the transition from data analysis to publication.

## Declarations

### Ethics approval and consent to participate

N/A

### Consent for publication

N/A

### Availability of data and materials

ENCODE datasets were downloaded from the ENCODE portal (https://www.encodeproject.org/) under accessions ENCFF144MRB, ENCFF405ZDL, ENCFF961SPZ, ENCFF188SZS, ENCFF118PBQ, ENCFF470HOG, ENCFF665RDD, and ENCFF727HQD.

### Authors’ contributions

NVB conceived the project under guidance from DCP. NVB wrote trackDJ’s code, performed analysis, and prepared figures. NVB and DCP wrote the manuscript.

## Acknowledgements

We thank all Page Lab members who tested the features of *trackDJ* and provided valuable feedback.

## Availability and Requirements

**Project name:** trackDJ

**Project home page**: https://github.com/neha-bokil/trackDJ

**Operating system(s):** Platform independent

**Programming language:** R

Other Requirements: R version 4.5 or higher, Bioconductor version 3.22 or higher

**License:** MIT

**Any restrictions to use by non-academics:** None

