## Supplemental Table and Figures for "Track Display Jockey (trackDJ): a user-friendly R package for visualization of epigenomic data"

| **ENCODE File** | **Assay** | **Track Type** | **Organism** | **File name** |
| --- | --- | --- | --- | --- |
| ENCFF144MRB.bigWig | H3K4me3 ChIP-seq | Coverage | Human | file1.bw |
| ENCFF405ZDL.bigWig | H3K4me3 ChIP-seq | Coverage | Human | file2.bw |
| ENCFF961SPZ.bed | H3K4me3 ChIP-seq | Peaks | Human | file3.bed |
| ENCFF188SZS.bed | H3K4me3 ChIP-seq | Peaks | Human | file4.bed |
| ENCFF511QFN.bedpe | CTCF ChIA-PET | Loops | Human | file5.bedpe |
| ENCFF470HOG.bigWig | H3K27me3 ChIP-seq | Coverage | Human | file6.bw |
| ENCFF665RDD.bigWig | H3K27me3 ChIP-seq | Coverage | Human | file7.bw |
| ENCFF727HQD.bigWig | H3K27me3 ChIP-seq | Coverage | Mouse | file8.bw |

Supplementary Table S1: Eight different ENCODE files were used in this study, along with their corresponding assays, track type, organism, and file name used in example code.

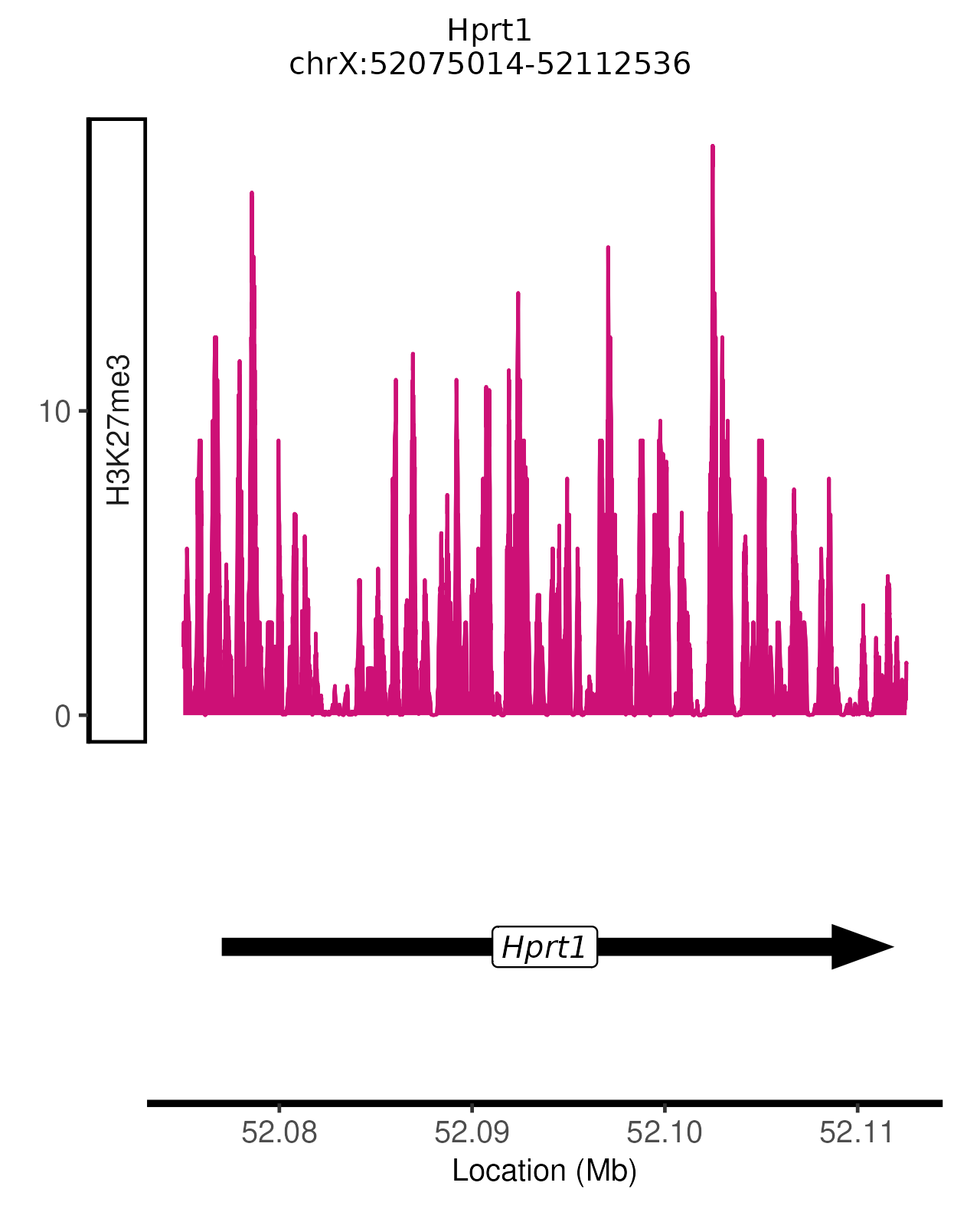

Figure S1: *trackDJ* plot of H3K27me3 at the *Hprt1* gene from the mmusculus_gene_ensembl annotation.

plot_genomic_tracks(genomicLoc=”Hprt1”, ensembl_set= ”mmusculus_gene_ensembl”, gene_symbol=”mgi_symbol”, covFiles=”file8.bw” covTrackColors=”deeppink3”, covTrackNames=”H3K27me3”)

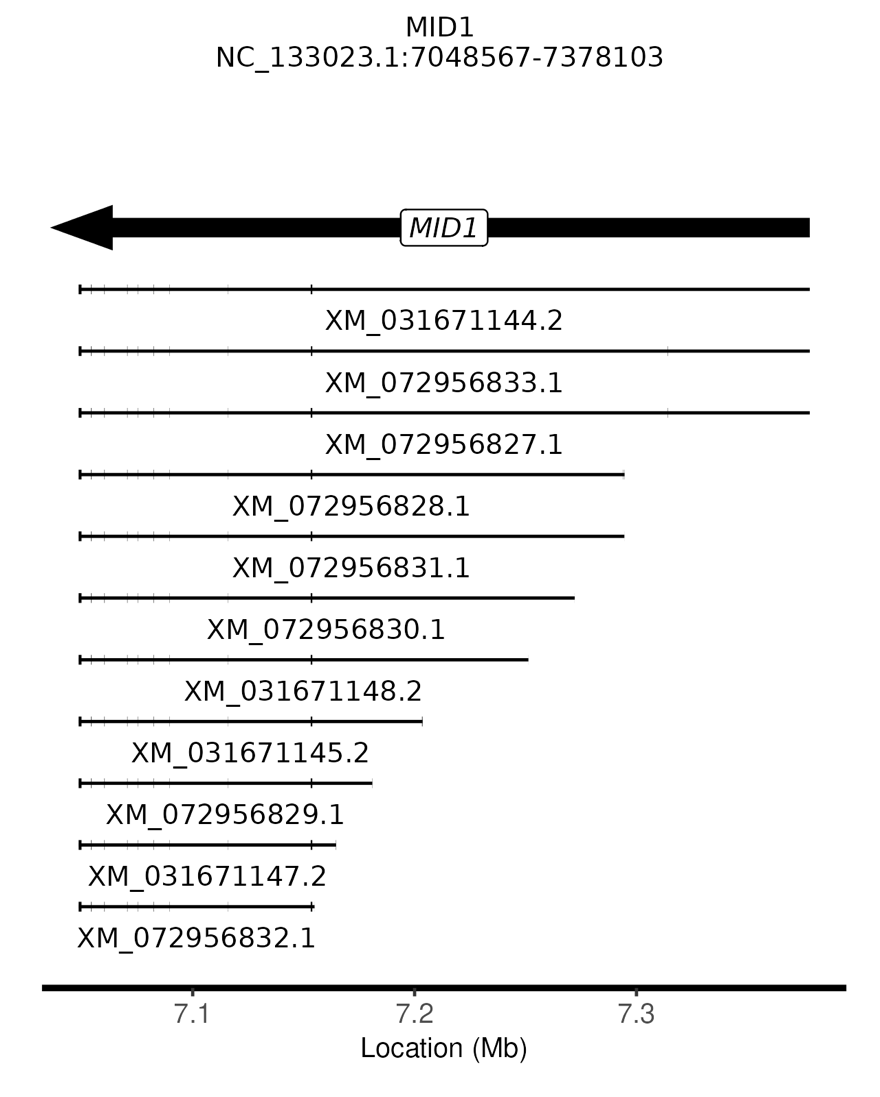

Figure S2: *trackDJ* plot of *MID1* using a gtf annotation file of the vicuña genome. The VicPac4 annotation file was downloaded from NCBI.

plot_genomic_tracks(genomicLoc=”MID1”, custom_anno=”VicPac4.gtf”, gene_symbol=”gene_id”, includeTranscripts=TRUE)

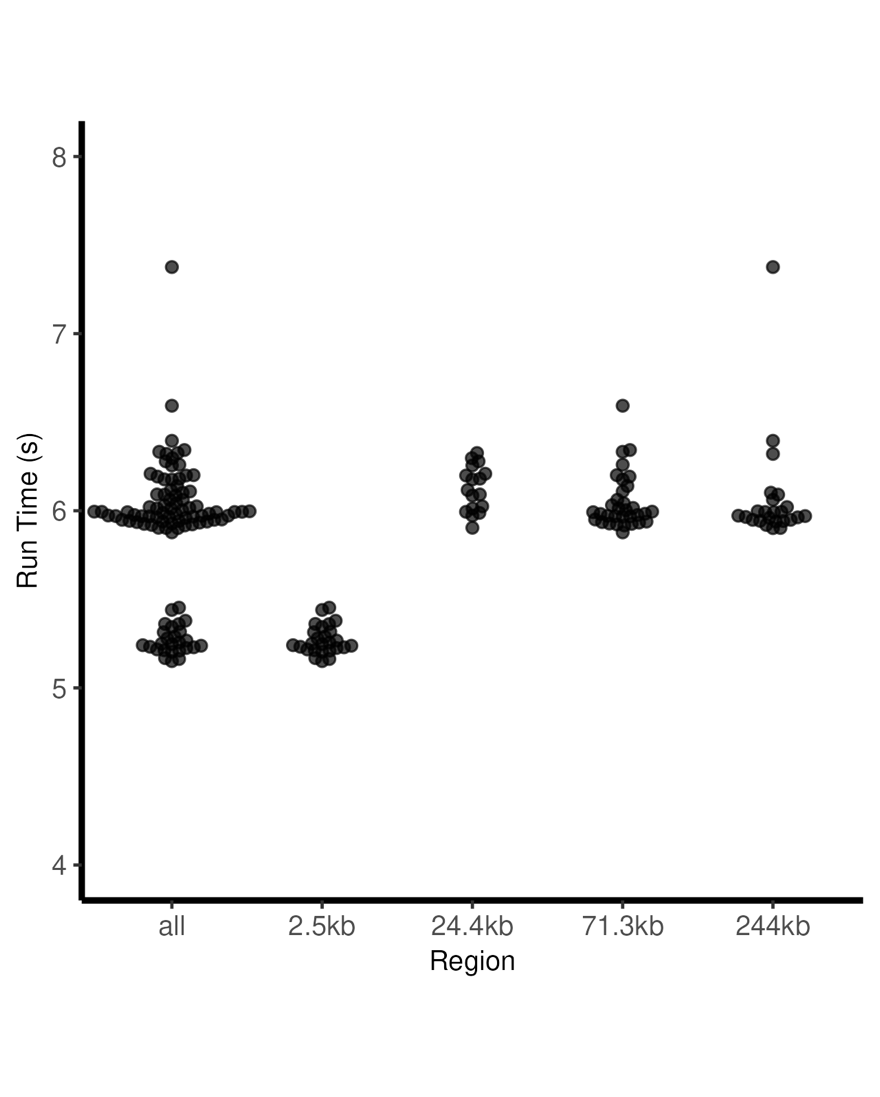

Figure S3: *trackDJ* runtimes range between 5-8 seconds for regions 2.5kb-250kb long. We ran *trackDJ* 100 times, each time randomly selecting one out of four possible regions of different lengths.
